# Viral susceptibility and innate immune competency of *Carollia perspicillata* bat cells produced for virological studies

**DOI:** 10.1101/2024.11.19.624190

**Authors:** Victoria Gonzalez, Cierra Word, Nahomi Guerra-Pilaquinga, Mitra Mazinani, Stephen Fawcett, Christine Portfors, Darryl Falzarano, Alison M. Kell, Rohit K. Jangra, Arinjay Banerjee, Stephanie N. Seifert, Michael Letko

**Affiliations:** Vaccine and Infectious Disease Organization (VIDO), University of Saskatchewan, Saskatoon, SK S7N 5E3 Canada; Department of Veterinary Microbiology, University of Saskatchewan, Saskatoon, SK S7N 5B4, Canada; Louisiana State University Health Sciences Center-Shreveport, Shreveport, Louisiana. 71103, USA; Paul G. Allen School for Global Health, Washington State University, Pullman, Washington, 99163, USA; Washington State University, Vancouver, Washington, 98686 USA; Department of Biology, University of Waterloo, Waterloo, ON N2L 3G1, Canada; Department of Laboratory Medicine and Pathobiology, University of Toronto, Toronto, ON M5S 1A8, Canada; Department of Molecular Genetics and Microbiology, University of New Mexico, Albuquerque, NM, 87131, USA; Department of Biochemistry and Molecular Biology, Faculty of Medicine, University of British Columbia, Vancouver, BC V6T 1Z3, Canada

**Keywords:** bat, innate immunology, virology, cell entry, virus-host interaction, interferon

## Abstract

Multiple viruses that are highly pathogenic in humans are known to have evolved in bats. How bats tolerate infection with these viruses, however, is poorly understood. As viruses engage in a wide range of interactions with their hosts, it is essential to study bat viruses in a system that resembles their natural environment like bat-derived *in vitro* cellular models. However, stable and accessible bat cell lines are not widely available for the broader scientific community. Here, we generated *in vitro* reagents for the Seba’s short-tailed bat (*Carollia perspicillata*), tested multiple methods of immortalization, and characterized their susceptibility to virus infection and response to immune stimulation. Using a pseudotyped virus library and authentic virus infections, we show that these *C. perspicillata* cell lines derived from a diverse array of tissues are susceptible to viruses bearing the glycoprotein of numerous orthohantaviruses, including Andes and Hantaan virus and are also susceptible to live hantavirus infection. Furthermore, stimulation with synthetic double-stranded RNA prior to infection with VSV and MERS-CoV induced a protective antiviral response, demonstrating the suitability of our cell lines to study the bat antiviral immune response. Taken together, the approaches outlined here will inform future efforts to develop *in vitro* tools for virology from non-model organisms and these *C. perspicillata* cell lines will enable studies on virus-host interactions in bats.

## INTRODUCTION

Most viral outbreaks in the last century have originated from cross-species transmission of viruses from wildlife to humans. For example, influenza viruses can spread to humans from birds and pigs, human immunodeficiency virus originated in chimpanzees, hantaviruses spillover from rodents, and mosquitos are essential vectors for several tropical arboviruses, including West Nile and Zika viruses. Decades of wildlife surveillance efforts have revealed that bats can carry a wide range of viruses. These include known human pathogens such as rabies and Marburg viruses, as well as sarbecoviruses (subgenus of coronavirus) found across Asia, Africa and Europe, and orthohantaviruses [1, 2]. Since 2000, the spillover of bat viruses into the human population has resulted in multiple, severe viral outbreaks, including SARS-CoV-2 and the resulting COVID-19 pandemic.

Despite the increasing research interests in bats and the pathogens they carry, there is a significant lack of laboratory tools for these animals. One confounding factor is the enormous diversity of bats, with over 1480 unique species spread across six continents [3]. Most of these species are understudied, leaving large knowledge gaps regarding how viruses transmit between bats and the animals they encounter. While some viruses are known to be carried by individual bat species, others are frequently shared between taxa [4]. Furthermore, comparative analyses show both variable and conserved components of bat immune pathways [5], which may suggest that bat-associated viruses may have evolved many mechanisms to evade the host immune response.

Another challenge to developing bat laboratory resources includes the species-specific interactions that often occur between viruses and their hosts. While some viruses may be able to replicate in multiple species, most viruses, including those found in bats, are species-specific and do not replicate efficiently or at all with commonly available laboratory reagents. For instance, Hendra virus was observed to have higher infection efficiency in cells derived from its natural reservoir, the black flying fox (*Pteropus alecto*), compared to Nipah virus which is primarily found in the large flying fox (*Pteropus vampyrus*) [6]. Additionally, Ebolavirus and Marburg virus have similar replication kinetics in cells derived from the Egyptian fruit bat (*Rousettus aegyptiacus*), the natural host for Marburg virus [7]. Meanwhile, modelling of Ebolavirus infection in the Jamaican fruit bat (*Artibeus jamaicensis*) demonstrated systemic infection and oral shedding of the virus, but this was not observed for Marburg virus [7]. These studies emphasize the role of innate host factors and varied immune responses triggered in different bat species, where viruses may exploit or antagonize these responses in the bat species they have co-evolved with. Thus, to truly study how bat viruses interact with their natural reservoir hosts, it is essential to study these viruses in experimental systems that closely mimic their natural environments.

Development of antiviral drugs and therapeutics requires an intricate understanding of viral biology, which can only be derived from laboratory experiments in either animal- or cell culture-based models. The first bat cell line was derived from the lungs of an adult female Mexican free-tailed bat (*Tadarida brasiliensis*) in 1965 by Kiazeff and colleagues and was made available through the ATCC cell line collection that year [8]. In the 50 years since then, only 3 additional bat cell lines have been made publicly available to researchers (R05T, R06E, EfK3; **supplemental table 1**). As researchers discover novel viruses in bats, it is imperative that laboratory tools accommodate the growing need to study these viruses and their spillover potential.

Here, we developed new bat cell lines and laboratory reagents from a diverse panel of tissues from the Seba’s short-tailed bat (*Carollia perspicillata*). These cell lines were screened for their ability to support infection by a diverse panel of viruses and further developed for routine mammalian cell culture and experiments to study bat viruses and innate immune responses. Our studies demonstrate that the resulting *C. perspicillata* cell lines are immunocompetent and mount a protective antiviral response upon double-stranded RNA (dsRNA) stimulation, making them suitable for future studies of bat-virus interactions and bat immunity. In addition, the kidney-derived cell lines are susceptible to a range of viruses, including Middle Eastern respiratory syndrome coronavirus (MERS-CoV), vesicular stomatitis virus (VSV), and orthohantaviruses. Notably, these cells support Andes orthohantavirus replication, which is closely related to the other orthohantaviruses naturally found in these bats [9, 10].

## RESULTS

### *Carollia perspicillata* primary cultures are susceptible to entry by diverse viruses

We set out to develop bat-derived cell lines specifically for virological research with several criteria for the resulting product: (1) the cells should originate from a species that can be bred in captivity and studied under laboratory conditions, (2) the genome sequence for the species should be available, (3) the cells should grow under standard mammalian cell culture conditions to ensure accessibility, (4) the cells should have an intact innate immune response in order to gain insight into virus-host interactions, and (5) for virological relevance, the cells should support infection with viruses that are native to the bat species.

With the first and second points of our selective criteria in mind, we obtained tissues collected from a single, 3-year-old male *C. perspicillata* (*Cp*) bat from a disease-free captive colony housed in the Washington State University campus in Vancouver, Washington (**Fig 1A**). Brain, lung, kidney, liver and spleen tissues were frozen until processing and primary cells were isolated following standard protease digestion and culturing techniques (see methods)[6, 11–14]. We then generated a panel of diverse viral glycoproteins representative of different viral families, entry pathways, and zoonotic origin. Additional focus was given to orthohanatviruses as close relatives of Andes orthohantavirus have been identified in *C. perspicillata* [9, 10, 15]. Primary *Cp* cells were susceptible to entry by many of the tested pseudoviruses, with the hantavirus and VSV glycoproteins mediating the strongest entry (**Fig 1B**). Notably, viral entry observed in primary cells was similar in intensity to several standard laboratory cell lines including Huh-7, Vero E6 and HEK-293T cells transduced to express coronavirus receptors, human dipeptidyl peptidase 4 (hDPP4) and human angiotensin converting enzyme 2 (hACE2) (**Fig 1C**), as well as an immortalized Jamaican fruit bat (*Artibeus jamaicensis*) cell line we developed previously (**Supplemental Fig 1A**)[16]. In contrast, several other bat cell lines that we developed in earlier studies, including immortalized kidney cells from the big brown bat (*Eptesicus fuscus*) and the Egyptian fruit bat (*Rousettus aegyptiacus*)[14, 17], were less susceptible to entry using our pseudotyped virus library (**Supplemental Fig 1B-D**).

**Figure 1.**
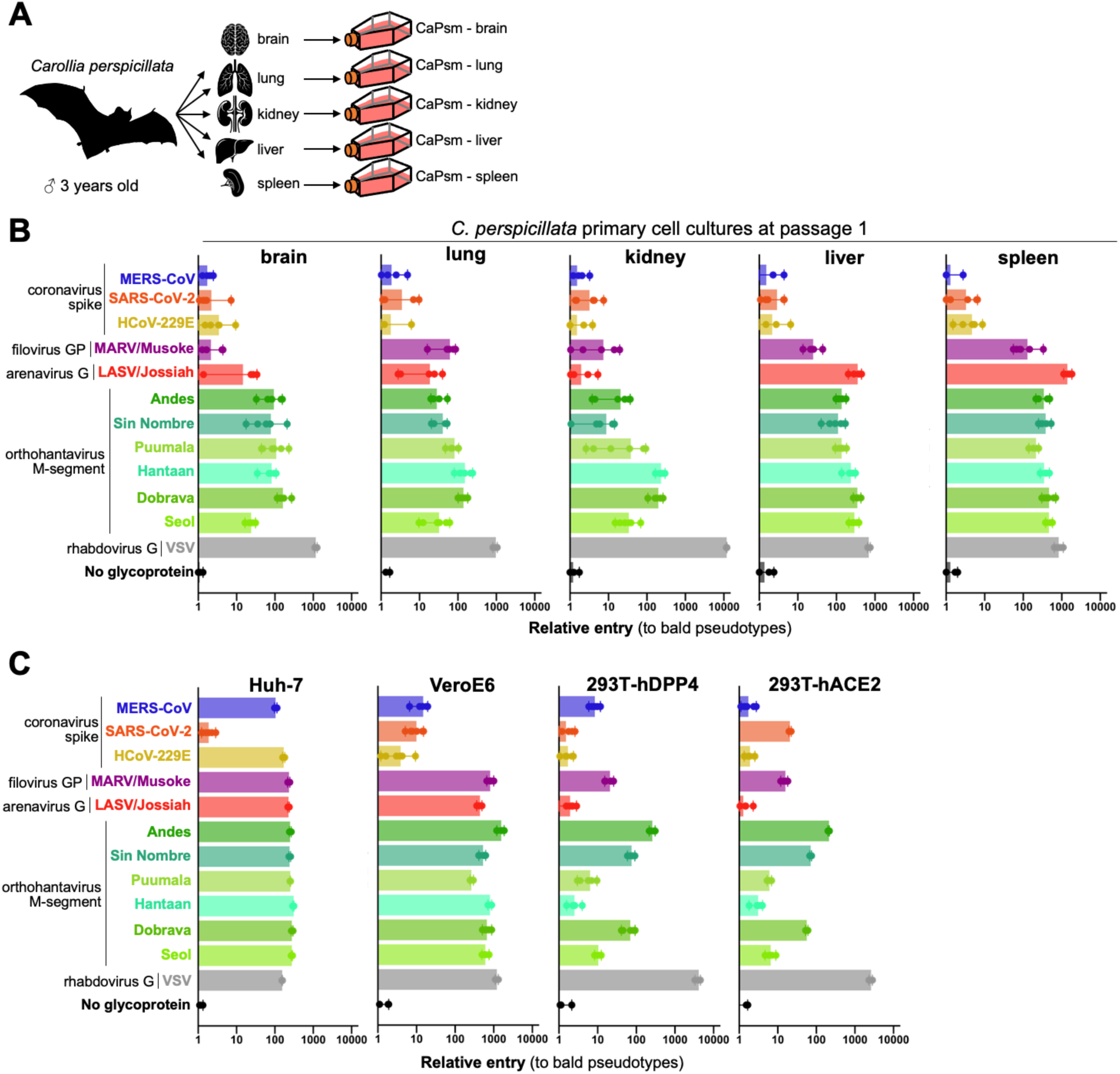
*Carollia perspicillata* primary cultures are susceptible to entry of diverse viruses. **(A)** Primary cells were isolated from brain, lung, kidney, liver and spleen of a single *C. perspicillata* bat and **(B)** infected with pseudotypes bearing glycoproteins from the indicated viruses. **(C)** Cell entry phenotypes of common laboratory cell lines to viral entry with pseudotype panel. Shown are results for 6 infection replicates per virus.

### Immortalization of *C. perspicillata* cell lines through diverse methods

While primary cells more faithfully mimic tissue-specific gene expression profiles, they are limited in their number of cell divisions and passages and are not a homogenous population. Not surprisingly, the majority of molecular virology studies are performed in immortalized cell lines that divide indefinitely and can therefore be easily shared among labs. Several methods have been used by various groups to immortalize primary cells. Viral proteins, such as the large T-antigen from SV40, inhibit tumor suppressor proteins like TP53 and Rab and shut down inhibition of cell division that normally follows in later passage primary cells [18, 19]. Similarly, the *Myotis polyomavirus* T antigen (MyPVT) has been shown to enhance DNA replication in Vero cells and immortalize *E. fuscus* kidney cells, though the mechanisms of action are unclear [14]. Over-expression of human telomerase reduces the genomic damage from continual cell divisions and has been used to immortalize bat cells previously [6, 20]. Directed knockout of the tumor suppressor, *TP53*, is yet another strategy to develop life-extended cells that has been employed more recently with the development of Cas9-targeted gene modification [21–24].

We tested all four strategies with multiple primary cell cultures using either in-house generated or commercially obtained lentivirus transduction systems (see methods). Interestingly, while some methods seemed to work for all the cell types tested, some cells stopped dividing after lentiviral transduction and were not further characterized (**Fig 2**). As we and others have noted, most of the immortalized cells were less susceptible to viral entry than their parental primary cells (**Fig 2B-E**) [6]. A few notable exceptions to this trend included *Carollia* brain cells transduced with TP53-guided Cas9 (gRNA: TP53), which showed high susceptibility to orthohantaviruses (**Fig 2B**), and kidney cells transduced with TP53-guided Cas9, which showed measurable susceptibility to MERS-CoV (**Fig 2C**). To test whether the cells retained their viral susceptibility over time, we performed entry assays on cells that were maintained for up to 31 passages (**Supplemental Fig 2**). Unfortunately, many of the transduced cells did not continually divide suggesting the approaches taken did not lead to immortalization. Three out of four of the immortalized cells evaluated broadly lost susceptibility to pseudovirus entry, with only the TP53-treated kidney cell line exhibiting an increase in susceptibility to MERS-CoV spike pseudotyped virus **(Supplemental Fig 2**).

**Figure 2.**
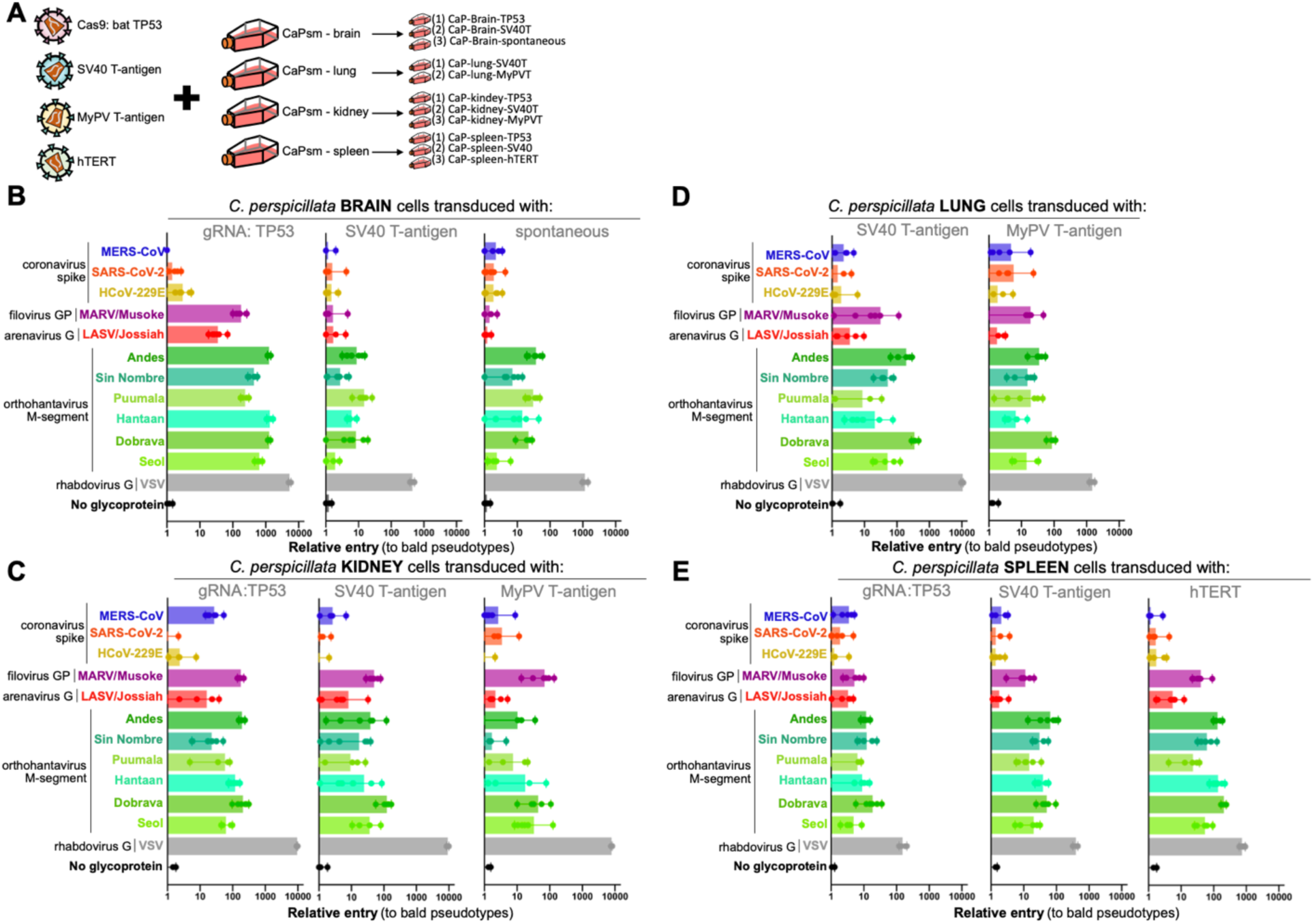
*Carollia perspicillata* cell lines immortalized through diverse methods are susceptible to viral entry. **(A)** Select primary cells were further transduced with lentiviral vectors representative of different strategies to immortalize cell lines. Transduced **(B)** brain, **(C)** lung, **(D)** kidney, and **(E)** spleen cells were screened for entry with pseudotyped particles bearing glycoproteins from the indicated viruses. Shown are the data from infection experiments perfomed in in 6-plo.

### *C. perspicillata* cells support infection with replication-competent recombinant VSV (rVSV)

To further assess if these primary cells were suitable for viral entry experiments, we tested a subset of primary cells for their ability to support infection with replication competent recombinant VSV (rVSV). VSV bearing its native glycoprotein or glycoproteins from Andes and Hantaan virus efficiently replicated in primary kidney cells (**Fig 3A**). To assess if Andes virus entry in these cells is mediated by a known hantavirus receptor, Protocadherin-1 (PCDH1), we incubated rVSVs with soluble PCDH1 (soluble extracellular cadherin domains 1 and 2, sEC1-2) and measured infection in *Cp* cells and human osteosarcoma U2OS cells, with the latter used as a positive control. Similarly to U2OS cells, a reduction in Andes virus glycoprotein-mediated infection was observed in *Cp* cells (**Fig 3B-C**). Notably, Hantaan virus was not inhibited in either cell line, in agreement with previous work showing that this virus does not use PCDH1 as a receptor [25]. However, preincubation of cells with a human PCDH1-specific antibody (mAb-3305) that has been shown to block PCDH1-dependent infection of Andes virus previously [25], did not block Andes virus glycoprotein-mediated infection in *Cp* cells but did so in human pulmonary microvascular endothelial cells (HPMEC) (**Fig 3D**). To determine if this was due to the rapid recycling of mAb-3305, the PCDH1-specific antibody was preincubated with *Cp* cells on ice (**Fig 3E**). Preincubation of the *Cp* cells with mAb-3305 did not block infection, suggesting that this antibody may not be cross-reactive or alternatively, Andes virus is capable of using an alternate entry receptor in these bat cells.

**Figure 3.**
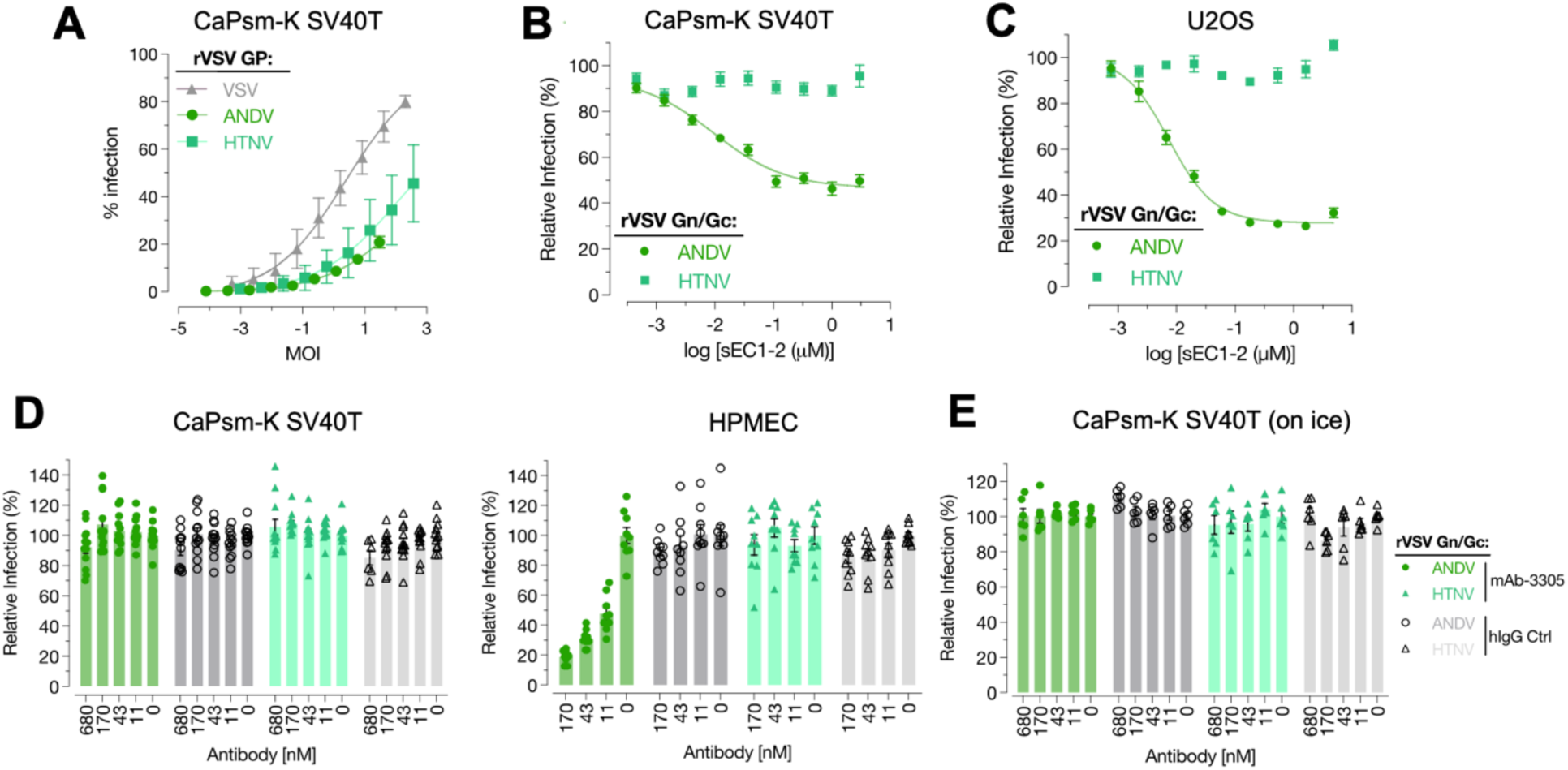
*C. perspicillata* cells support infection with replication-competent recombinant VSV (rVSV). **(A)** *Cp* kidney cells immortalized by SV40 T-antigen (CaPsm-K SV40T) were infected with replication competent VSV (rVSV) pseudotyped with indicated orthohantavirus glycoproteins. Neutralization of rVSV ANDV and HTNV with soluble PCDH1 receptor (sEC1-2) on *Cp* kidney cells **(B)**, and human U2OS cells for comparison **(C)**. **(D) N**eutralization of rVSV-ANDV and HTNV on *Cp* kidney cells by preincubation of cells with an anti-PCDH1 (mAb-3305) antibody. **(E)** Neutralization of rVSV ANDV by mAb-3305 on human pulmonary microvascular endothelial cells (HPMEC) is shown as a positive control. To rule out the possibility that mAb-3305 fails to block rVSV-ANDV infection in *Cp* kidney cells due to rapid uptake/clearance of the antibody, the experiment was performed on ice.

### Immortalized cells respond to poly(I:C) stimulation

To test the immunocompetence of the *C. perspicillata* cell lines, we first measured the transfectability of both primary and immortalized kidney cell cultures using poly(I:C) (surrogate viral dsRNA) labelled with rhodamine (poly(I:C)-Rho) (**Fig 4A**). Poly(I:C)-Rho was successfully delivered in all cell lines, with the TP53-guided cell line demonstrating the highest uptake of the immunostimulant (**Fig. 4A**). Next, we characterized the response of these bat cells to poly(I:C) by measuring interferon beta (*IFN*β) and interferon induced protein with tetratricopeptide repeats 1 (*IFIT1*) transcript levels using qPCR (**Fig 4B**). All four cell lines upregulated *IFN*β and *IFIT1* transcript levels to similar levels. To assess whether the IFNβ response was effective in blocking virus infection, we stimulated cells with poly(I:C) prior to infection with VSV that was engineered to express the green fluorescent protein (VSV-GFP) (**Fig 4C-D**). While VSV-GFP replicated efficiently in both primary and immortalized kidney cells, TP53-guided cells were the least permissive. However, pretreatment with poly(I:C) completely inhibited virus replication. Immunoblotting for IFIT1, an interferon stimulated gene (ISG), confirmed that poly(I:C) induced the expression of IFIT1 in infected and uninfected cells (**Fig 4E**).

**Figure 4.**
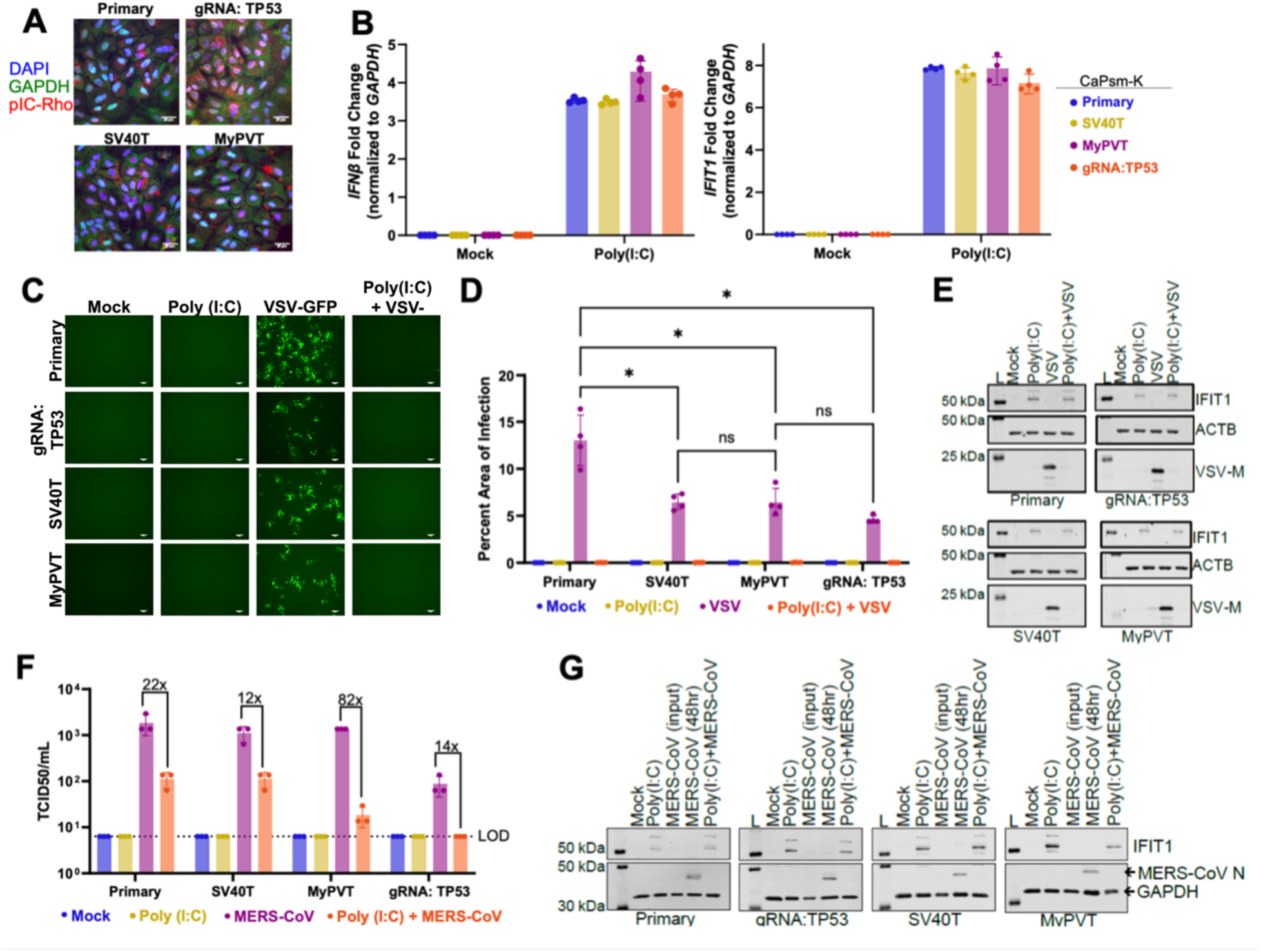
Immortalized *Carollia* kidney cells respond to surrogate virus infection. **(A)** CaPsm-K primary and immortalized cells were transfected with 10 μg of poly(I:C) chemically labelled with rhodamine (pIC-Rho) for 16 hours. Following fixation, cells were stained with antibodies against GAPDH and DAPI and visualized by confocal microscopy. Scale bars represent 25 μm. **(B)** CaPsm-K cells were transfected with 100ng of poly(I:C) for 8 hours. The upregulation of *IFN-β* and *IFIT1* transcripts was assessed by qPCR. Data are represented as a mean ± SD, n=4 replicates. **(C)** CaPsm-K cells transfected with 100 ng of poly(I:C) for 8 hours were infected with VSV-GFP (MOI 1) for 16 hours (n=4). Viral replication was visualized by fluorescent microscopy. Scale bars represent 100 μm. **(D)** GFP signal was quantified using ImageJ (Two-way ANOVA with Šídák’s multiple comparisons test). **(E)** Representative western blot of VSV-GFP infected cells, where IFIT1, VSV-M and ACTB were probed for. L = molecular weight ladder. **(F)** CaPsm-K cells transfected with 100 ng of poly(I:C) for 8 hours were infected with MERS-CoV (MOI 0.1) for 48 hours (n=3). Supernatant was collected and TCID50 assay was performed to assess viral titer. **(G)** Representative western blot of MERS-CoV infected cells, where IFIT1, MERS-CoV nucleocapsid (N), and GAPDH were probed for. L = molecular weight ladder.

Our entry results using pseudotyped virus suggested that the *Cp* kidney cell lines expressed the necessary factors for MERS-CoV entry (**Fig 2C**). Therefore, we repeated our poly(I:C) experiment but replaced VSV with authentic MERS-CoV (**Fig 4F**). Similar to VSV-GFP, all cell lines were permissive to infection with MERS-CoV, with the virus replicating to lower titers in TP53-guided cells. Stimulation with poly(I:C) prior to infection reduced MERS-CoV replication, with the most substantial reduction observed in kidney cells immortalized using MyPV GP5. IFIT1 analysis through immunoblotting demonstrated similar expression of IFIT1 in cells pre-stimulated with poly(I:C), with IFIT1 not induced in cells only infected with MERS-CoV (**Fig 4G**). Taken together, these results demonstrate that *C. perscipillata* kidney cells are suitable for both virological and immunological studies; however, differences in virus susceptibility exist and should be considered in future studies.

### Immortalized clonal cell lines retain susceptibility to viral entry

Our entry results suggest that the *C. perspicillata* kidney cell lines transduced with TP53-guided Cas9 are susceptible to entry via the widest range of viral glycoproteins of all the cell lines developed (**Fig 2C, Supplemental Fig 2**). Importantly, despite immortalization, these cells also retained their interferon response (**Fig 4**). We observed that later passage *Cp* TP53-guided cells were more susceptible to MERS-CoV than earlier passages (**Supplemental Fig 2**), and that this entry phenotype was more variable throughout the course of this study. Therefore, we wondered whether the *Cp* cells represented a mixed population of susceptible and non-susceptible cells. To help address this question, we isolated individual cells through limiting dilution plating (**Fig 5A**; see methods). Twenty clones were selected from the limiting dilutions for their ability to form dense monolayers. From these 20 clones, only five retained the ability to divide from one cell through plating for infection. Testing the viral pseudotype panel on these cells revealed minor variation in viral susceptibility between clones and an overall improvement in entry from the heterogenous parental population (**Fig 5B**). Of the five clones that were tested, only a subset continuously grew in culture, with clone-9 (CKg9) exhibiting the best growth and highest susceptibility to viral entry through continuous passage (**Fig 5C**).

**Figure 5.**
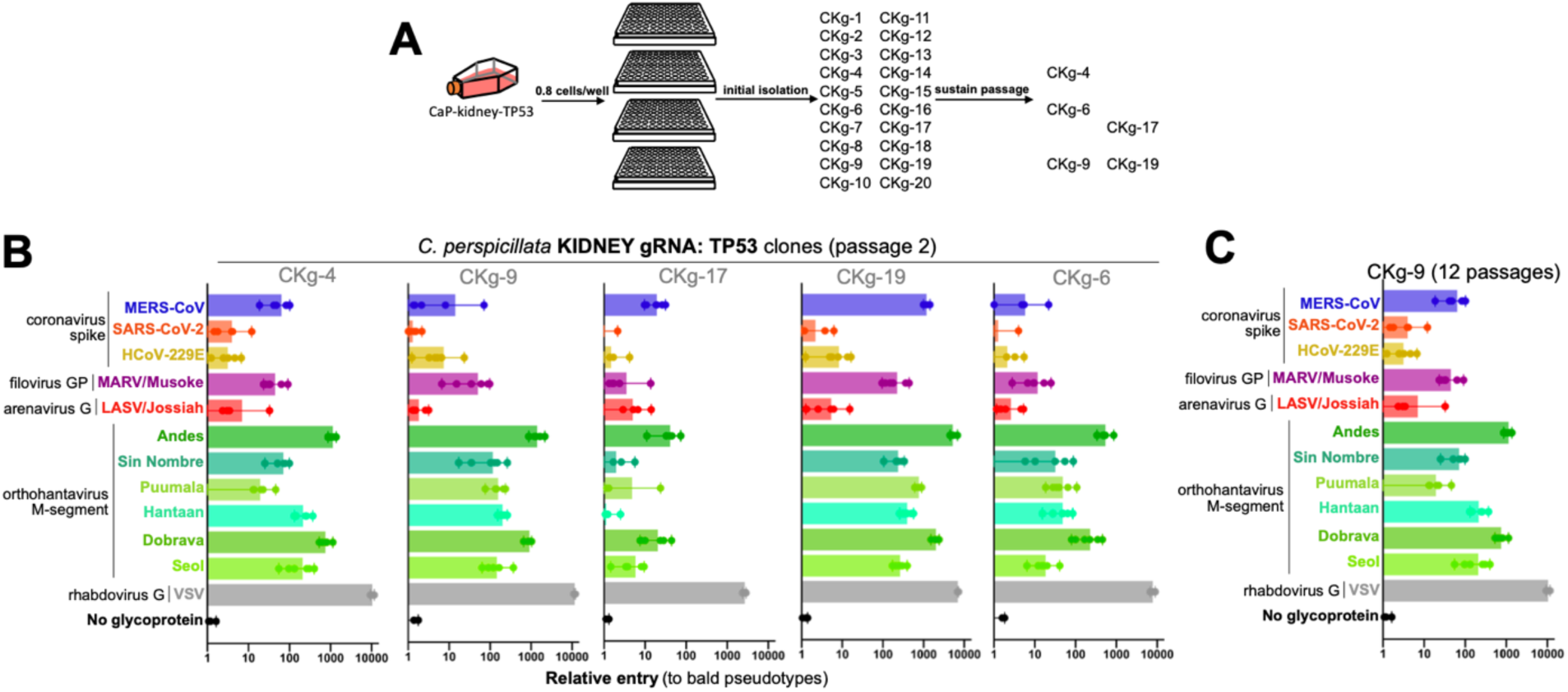
Immortalized clonal cell lines retain susceptibility to viral entry. **(A)** Select transduced cells were further plated for clonal isolation and 20 clones were propagated from single cells plated in 96-well format. **(B)** Clones that continuously grew were further screened for suppressibility to viral entry with pseudotypes bearing glycoproteins of the indicated viruses. **(C)** Clones that grew past 10 passages were re-screened for susceptibility to viral entry. Shown are the results from infection experiments performed in 6-plo.

### Immortalized *Carollia* cells support whole virus replication

To further assess the suitability of these cells for virological experiments, we infected primary and immortalized *Cp* kidney cells with authentic VSV-GFP (MOI 1), MERS-CoV (MOI 0.1), and SARS-CoV-2 (MOI 0.01) for 1 to 72 hours (**Fig 6A**). Upon infection with VSV-GFP, all cell lines supported virus replication, with viral titers peaking at 24 hours in the primary cells. However, VSV-GFP titers did not peak until 48 hours in SV40T and MyPV GP5 cells, and until 72 hours in TP53-guided cells. Similarly, TP53-guided cells were the least permissive to MERS-CoV, with titers remaining below 10^2^ TCID_50_/ml by 72 hours of infection (**Fig 6A**). Meanwhile, similar replication kinetics were observed for infected primary and SV40T-immortalized cells, with MERS-CoV titers peaking around 10^3^ TCID_50_/ml by 72 hours of infection. For cells immortalized using MyPV GP5, infection peaked and plateaued at 48 hours. Consistent with our previous data which demonstrated that the *Cp* kidney cells do not permit entry of pseudoviruses displaying the SARS-CoV-2 spike protein (**Fig 2C**), infection with authentic SARS-CoV-2 was not supported in neither the primary nor immortalized kidney cell lines (**Fig 6A**).

**Figure 6.**
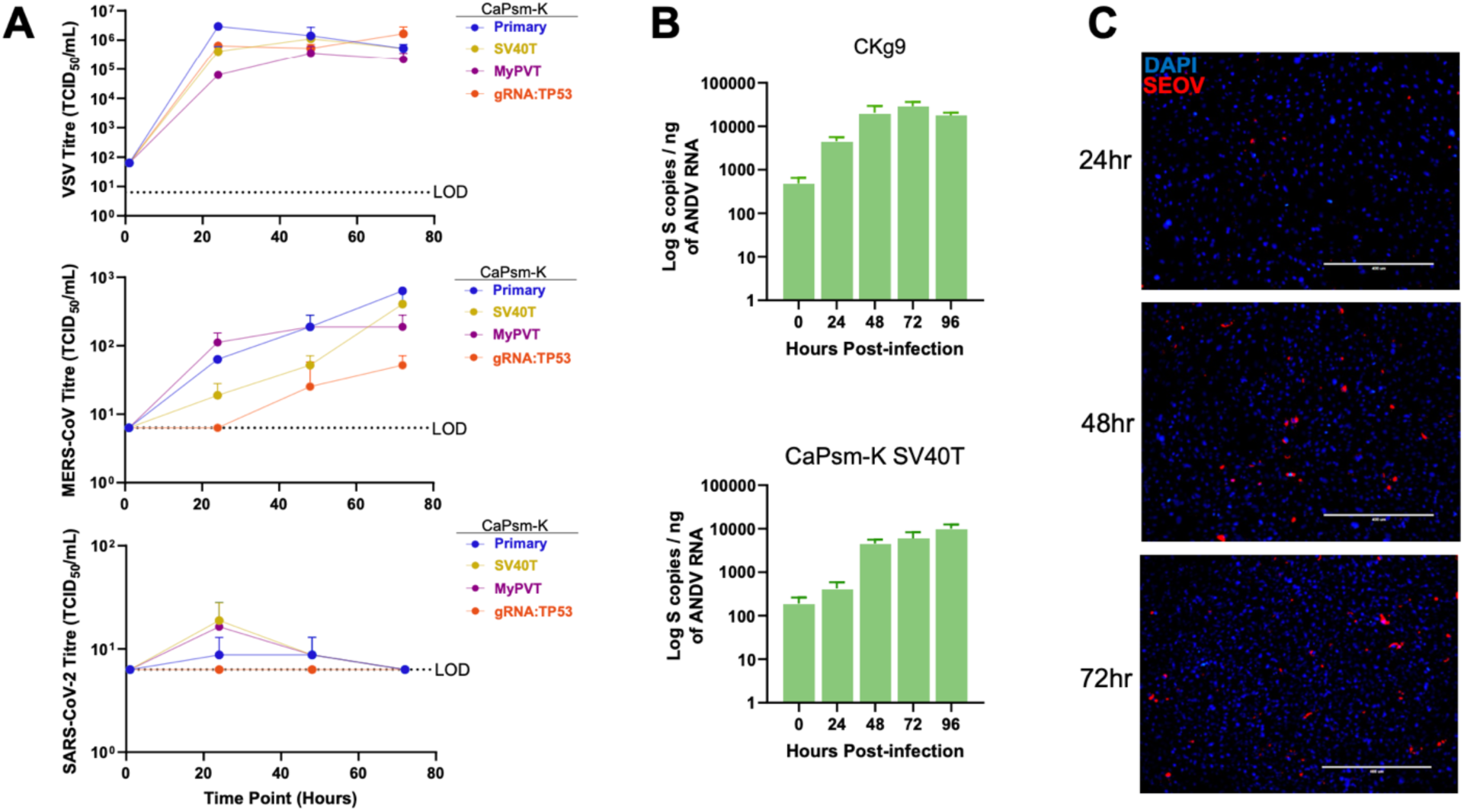
Immortalized *Carollia* cells support whole virus replication. **(A)** Primary and immortalized kidney (CaPsm-K) cells were infected with VSV-GFP (MOI 1), MERS-CoV (MOI 0.1), or SARS-CoV-2 (MOI 0.01) for up to 72 hours. Replication was monitored using TCID_50_ limiting dilution assays. **(B)** Clonal transduced kidney cells (CKg9) cells and CaPsmK-SV40T cells infected with Andes virus were monitored by qPCR. **(C)** CKg9 cells infected with Seoul virus (SEOV) were monitored by end-point microscopy for Seoul nucleocapsid protein.

As orthohantaviruses related to Andes virus have been reported in *C. perspicillata* [9, 10, 15], we subsequently tested the infectivity of Andes virus in the clonal transduced Ckg9 cell line and observed day-over-day increases in the viral genome suggestive of viral replication (**Fig 6B**). Infection of these cells with Seoul virus, another rodent-borne orthohantavirus, allowed for detection of Seoul nucleocapsid protein in the cell lines (**Fig. 6C**), suggesting that the Ckg9 cell line supports Seoul virus replication.

## DISCUSSION

Some bat species have been known to carry viruses that can infect people since the early 1930’s, when Joseph Pawan first described transmission of rabies virus from the common vampire bat (*Desmodus rotundus*) to humans in Trinidad [26, 27]. Since these initial findings, outbreaks of bat-derived viruses in humans have become an unfortunate but more frequent occurrence. The COVID-19 pandemic and subsequent identification of several closely-related viruses in bats underscores the urgency to understand bats and the viruses they carry [28, 29].

Our knowledge of viruses derives from our ability to study them in the laboratory. Despite nearly a century of bat virus research, there are very few standardized tools available for researchers to study bat viruses. Bat-derived cell lines are essential to studying molecular interactions between bats and their viruses. While several bat cell lines have been developed by various groups, most of them are not publicly available, and the few that are do not support replication of most viruses [6, 11, 30–32]. For example, while Chinese rufous horseshoe bats (*Rhinolophus sinicus*) harbor viruses related to SARS-CoV, primary and immortalized cell cultures from these animals do not support SARS-CoV or SARS-CoV-2 infection [11, 30]. Bat cell lines that do support viral replication often remain in the lab that develops them. Without common laboratory reagents available to more researchers, the long-standing question of how bats tolerate viruses that are otherwise pathogenic in other animals may remain elusive. Here, we developed new bat cell lines, tested methods to immortalize and subclone them, introduced gene disruptions, and measured innate interferon responses. Importantly, because virological studies are the goal of our cell lines, we continually assessed their ability to support viral entry and replication throughout the study and discontinued the use of cells that lost this ability. The resulting cells grow under standard laboratory conditions with accessible and cost-effective cell culture media and support viral growth and innate immune responses.

Changes in cell gene expression are a major hurdle for developing cell lines. As mammalian cell cultures age, their expression profile changes over time [33]. Routine laboratory cell culture includes the use of fetal-bovine serum, which does not contain species-specific growth factors for bat cells, or other non-bovine species. Our results showed differences in viral susceptibility between primary, early- and late-passage immortalized cells (**Fig. 1, 2, S2**). As these cells were all derived from the same individual, and share similar interferon response profiles, this variation in cell phenotypes - in particular specific coronavirus entry profiles - suggests shifts in receptor expression, which has been documented for coronavirus receptors in other bat cell lines [30]. Nevertheless, some cell lines retained their susceptibility and additional screening of clonal immortalized derivatives allowed us to identify cell lines that retained susceptibility to coronaviruses (**Fig 2**). Thus, receptor downregulation appears stochastic and not necessarily a factor of cell culture age. Taken together, these findings should be taken into consideration when developing animal derived cell lines, with the inclusion of clonal screening to identify key subpopulations of cells with desired laboratory properties.

The *Carollia* cell lines developed in this study supported strong entry for all the pseudotyped orthohantaviruses tested in our assays and were capable of supporting infection with *bona fide* Andes and Seoul viruses (**Fig 1, 3, 6B-C**). The receptor for New World orthohantaviruses, which includes close relatives of orthohantaviruses found in *Carollia* bats, has been shown to be PCDH1 [25]. The soluble receptor (sEC1-2) experiments show that PCDH1 may mediate Andes virus entry in these cells. However, the mAb-3305 experiments suggest that either this antibody does not recognize bat PCDH1 due to sequence divergence or bat PCDH1 may not be the only receptor required for Andes virus entry in these bat cells (**Fig. 3**). Alternatively, sEC1-2 may have blocked Andes virus entry by steric hindrance and another unknown receptor mediates hantavirus entry in these cells. Recent work has also demonstrated the expansion of the Buritiense hantavirus amongst *Carollia* species that circulate near human communities in Brazil [34], increasing the risk of spillover. Indeed, our study and these cell lines will provide researchers with a unique opportunity to study hantaviruses in their bat hosts.

Bats are known to carry a wide range of viruses, some of which are highly pathogenic in humans or are closely related to such viruses. Remarkably, bats do not exhibit obvious signs of disease from this viral burden, with the unexplained exception of Tacaribe virus, Rabies virus, and Lloviu virus [35–37]. This tolerance to viral infection is a significant area of research, with the goal of one day adapting these mechanisms in the form of human therapies. Cell lines developed in this study will be helpful in understanding virus-host interactions in *C. perspicillata* bats using viruses that these bats are speculated to naturally harbor or be exposed to like orthohantaviruses and influenza A-like viruses (H18N11).

## METHODS

### Biosafety and Ethics Statement

Bat tissues were sourced from a US-based facility at Washington State University (Vancouver, WA, USA). Collection of animal tissues used in this study was approved by Washington State University IACUC (ASAF #6588). Experiments with live, replication competent coronaviruses were performed at the University of Saskatchewan/VIDO’s BSL3 facility following approved protocols and standard procedures. Experiments with live hantaviruses were performed under BSL3 conditions at the University of New Mexico. Experiments with replication defective VSV pseudotypes and replication competent rVSVs were performed at BSL2+ conditions at Washington State University and LSU Health Sciences Center-Shreveport, respectively.

### Isolation of primary cells

Tissues were harvested from a male *Carollia perspicillata* bat at approximately 20 months of age, 17.1 g, from a disease-free colony. Tissue samples were washed and finely minced in sterile phosphate buffered saline (PBS), transferred to fetal bovine serum (FBS) supplemented with 10% dimethyl sulfoxide and frozen in liquid nitrogen. Primary cells were isolated as we have previously reported for other bat species [13, 14]. Briefly, samples were thawed at 37°C and transferred to sterile 10cm plates. FBS/DMSO solution was removed with a pipette and tissues were washed with PBS. Tissues were incubated in trypsin for 10 minutes, with gentle agitation through a 1000uL pipette to help loosen cells from the tissue matrix. Trypsin was neutralized with primary cell culture medium consisting of high glucose DMEM (Corning), 12% FBS, penicillin/streptomycin, amphotericin B, L-glutamine, and minimum essential medium non-essential amino acids. The slurry of cells, tissue debris and media were transferred to 75cm^2^ flasks, labeled “passage 0” and maintained at 37°C with 5% CO_2_. Flasks were monitored daily for cell growth and media was refreshed every two days. Flasks that achieved 70-90% confluency with primary cell cultures were expanded into larger 150cm^2^ flasks for banking stocks of early passage cells and future experiments. Primary cells were used up to passage 10 for viral infection assays in Figure 4.

### Immortalization of primary cells

Immortalization of bat cells with the large T-antigen form SV-40 virus and the human telomerase gene (hTERT) have been previously described [6, 13]. Lentiviruses encoding these proteins and puromycin N-acetyletransferase were obtained commercially and used following manufacturers instructions (GenTarget). Cells were transduced with lentiviruses encoding either SV-40 T-antigen or hTERT and then maintained in a primary cell cuture media (see above) with puromycin at between 0.5-1.5ug/mL depending on the cell source for two passages. After bulk, untransduced cell die-off, the media was changed to a standard maintenance media consisting of high-glucose DMEM supplemented with 10% FBS, L-glutamine and penicillin streptomycin. MyPV is an ortholog of the SV-40 T-antigen but from a bat polyomavirus and has been described previously described [14]. Cells transduced with MyPV lentivirus were maintained in standard 10% FBS DMEM. TP53 KO was inspired by previous studies that used human, mouse and canine cells [22–24]. At the time of designing this study, the sequenced genome for *Carollia perspicillata* was still in development. Therefore, we used the *Rousettus aegyptiacus* (GenBank: (NW_023416309.1:73376916-73388351) and *Artibeus jamaicensis* (GenBank: (NW_026521968.1:7682714-7693614) genomes to design a series of Cas9-guide RNAs directed toward the exon-intron junctions of exons 1 and 5 of the bat TP53 tumor suppressor gene. Four resulting gRNAs were designed:

RA TP53 – EXON 1 – FORWARD: 5’-CTTGTGGAAACTGTGAGTAG-3’
RA TP53 – EXON 5 – REVERSE: 5’-TCCACCCGGATAAGATGCTG-3’
AJ TP53 – EXON 1 – REVERSE: 5’-TGACTCATTGGTGGCTCCAC-3’
AJ TP53 – EXON 5 – FORWARD: 5’-GGTGCCCTACGAAACACCTG-3’

Guide RNAs were cloned into a lentiviral Cas9/CRISPR vector (addgene #52961)[38] and used to produce VSV-g pseudotyped lentiviral particles in 293T cells. Cas9:bat TP53 lentiviruses were collected 48 hours later, filtered through 0.45 μM mixed cellulose ester filters and stored at - 80°C until use. Cells were transduced with a combination of all four lentiviruses and maintained in standard media consisting of high-glucose DMEM supplemented with 10% FBS, L-glutamine and penicillin/streptomycin. All transduced cells were expanded in large culture flasks for stock banking shortly after transduction and, when applicable, puromycin selection.

### Clonal cell lines

Early passage stocks of *Carollia perspicillata* kidney cells transduced with Cas9:bat TP53 lentivirus were thawed and maintained for one passage before seeding at low density of 0.8 cells/well in 96 well plates. Wells were monitored over two weeks for single cell isolation and growth. Twenty wells that reached 100% confluency the fastest were selected for expansion into 6-wells. Five of the 20 clones maintained continual growth to be expanded into larger culture flasks. Clonal cells were maintained in standard media consisting of high-glucose DMEM supplemented with 10% FBS, L-glutamine and penicillin/streptomycin. DMEM with high glucose (DMEM; Gibco, #12491015) supplemented with 10% fetal bovine serum (FBS; Sigma-Aldrich, #12107C) and 1X antibiotic antimycotic (Gibco, #15240062).

### Plasmids

Glycoproteins from MERS-CoV/EMC12, SARS-CoV-2/Wuh1, HCoV-229E, and Lassavirus/Josiah were codon optimized and cloned into pcDNA3.1 as previously described [13, 39, 40]. Marburg/Musoke glycoprotein in pCAGGS vector was generously provided by Andrea Marzi (NIH/NIAID). Orthohantavirus M-segments from plasmids generously provided by Jay Hooper (USAMRIID) were subcloned into pcDNA3.1 using standard PCR approaches. Plasmids used for the expression of His-tagged version of soluble extracellular domains 1-2 (sEC1-2) of human PCDH1 and mAb-3305, a human IgG targeting EC1 domain of human PCDH1, were described previously [9, 10, 15].

### Pseudotype entry assay

Pseudotype entry assays were performed as described previously [13, 39, 40]. Briefly, 293Ts were seeded in poly-lysine treated 6-well plates and transfected with plasmids encoding viral glycoproteins. Twenty-four hours later, cells were infected with single cycle vesicular stomatitis luciferase-GFP dual reporter virus pseudotyped with VSV-G (VSVΔG-luc/GFP) for 1 hour and washed three times with PBS to remove residual input virus. Culture supernatants containing viral psuedotpyes were collected 48 hours later, briefly centrifuged to remove debris, aliquoted and stored at −80°C. For entry assays, cells were seed in 96-well plates and infeected with equal volumes of viral pseudotypes diluted in DMEM supplemented with 2% FBS, l-glutamine and penicillin/streptomycin. Innoculated cells were centrifuged at 1200xG, 4°C for one hour and then transferred to 37°C incubators with 5% CO_2_. Luciferase was measured as a readout for cell entry (BrightGlo; Promega), and data were normalized by comparing raw luciferase signal from pseudotyped viruses against luciferase signal from non-pseudotyped negative control particles [13, 17, 41, 42].

### rVSV infection assays

Titers of the stocks of rVSVs expressing Andes virus (ANDV) Gn/Gc, Hantaan virus (HTNV) Gn/Gc or VSV G glycoproteins as their sole entry proteins were determined on Vero cells to determine multiplicity of infection (MOI) for infection experiments on *Cp* cells. Vero cells plated in 96-well plates (8,000 cells per well) were infected with serial log dilutions of rVSVs. At 8 hours post-infection, the cells were fixed using 2% formaldehyde for 15 minutes at room temperature and nuclei were counter-stained with Hoechst 33342. Infected (eGFP-positive) cells were imaged on a Cytation-5 (Agilent) and automatically enumerated using the onboard Gen5.0 software to calculate titers of the stocks. For comparison of infectivity of these rVSVs on *Cp* cells, immortalized kidney cells plated in 96 well plates (8,000 cells per well) were infected with 2-fold serial dilutions of viruses (an MOI range of 1.5 to 374 for rVSV Andes virus and 0.8 to 207 for rVSV Hantaan virus and VSV G). At 16 hours post-infection, cells were fixed and scored for infection as described above. The results were expressed as percent infection.

For neutralization of rVSV infection with sEC1-2, rVSV-ANDV Gn/Gc and HTNV Gn/Gc viruses were titered to determine the amount of virus required to obtain 20-30% infection as 16 hours post-infection in immortalized *Cp* kidney cells and human osterosarcoma U2OS cells. These pretitrated amounts were incubated with 3-fold serial dilutions of sEC1-2 starting at 3 μM for one hour at 37℃. These rVSV:sEC1-2 mixtures were then applied to monolayers of immortalized *Cp* kidney cells in 96-well plate and infection was scored at 16 hours post-infection as described above. The results were expressed as relative percent infection as compared to that in well without sEC1-2 protein (infection in no sEC1-2 well was set at 100%).

The rVSV neutralization experiments with of rVSV infection with an anti-PCDH1 antibody, mAb-3305 were performed similarly except that the cells were incubated with mAb-3305 or a human IgG control (Ctrl) at 11 to 680 nM concentration in 4-fold serial dilutions for one hour at 37℃ or on ice before infection with pre-titrated amounts of rVSV-ANDV Gn/Gc or HTNV Gn/Gc. For on ice experiments, cell monolayers were prechilled on ice for 10 minutes before the addition of antibodies. At 16 hours post-infection, the percent relative infection was estimated as described above as compared to that in the well without antibody (infection in no antibody wells was set at 100%). Primary human pulmonary microvascular cells (HPMECs) were used as positive controls for these studies.

### Viruses

Genetically engineered vesicular stomatitis virus encoding a green fluorescent protein (VSV-GFP) cassette were propagated in Vero76 cells in DMEM and stored at −80°C. Middle East respiratory syndrome coronavirus (MERS-CoV; isolate EMC/2012) and ancestral severe acute respiratory syndrome coronavirus 2 (SARS-CoV-2; isolate SB3) was propagated in Vero76 cells [43]. All work with infectious MERS-CoV and SARS-CoV-2 was completed at VIDO in a containment level 3 laboratory and was approved by the institutional biosafety committee. Standard operating procedures approved by the institutional biosafety committee were followed for sample inactivation.

Seoul virus strain SR11 and Andes virus strain CHI-9717869 were propagated on Vero E6 cells (ATCC, CRL-1586) for 12 days. Infectious virus was isolated by harvesting supernatant and centrifuging at 1000rcf for 10 minutes to remove cellular debris. The rVSVs expressing Andes virus and Hantaan virus glycoproteins have been described previously [44].

### Transfection

Cells were seeded at a density of 1.5×10^5^ cells/well for 24 hours followed by transfection with 10 ug of poly(I:C) chemically labeled with rhodamine (Invivogen, #tlrl-picr) for 16 hours, or with 100 ng of poly(I:C) (Invivogen, #tlrl-pic) for 8 hours. Cells transfected with poly(I:C) chemically labeled with rhodamine were fixed in methanol and stained for GAPDH (EMD Millipore, #MAB374) and DAPI (Thermo Scientific, #62248) and visualized using confocal microscopy.

### Whole-virus infection

*Carollia* kidney cells were seeded at a seeding density of 1.5×10^5^ cells/well. Following 24 hours, cells were transfected with 100 ng of poly(I:C) (Invivogen, #tlrl-pic) for 8 hours prior to infection with VSV-GFP (MOI 1) or MERS-CoV (MOI 0.1) for 16 and 48 hours, respectively. Protein lysate was harvested and assessed by immunoblotting, while supernatant for MERS-CoV infected cells was harvested to assess viral titer by tissue culture infectious dose assay (TCID_50_). Kidney cells were similarly seeded for VSV-GFP (MOI 1), MERS-CoV (MOI 0.1), and SARS-CoV-2 (MOI 0.01) growth curves. Supernatant was harvested following 1, 24, 48, and 72 hours of infection and titerd on Vero76 cells using a TCID_50_ assay. For Seoul and Andes virus infections, cells were seeded in cell culture vessels 18-24 hours prior to infection at a target density of 70%. Virus stock was diluted to the target, cell-specific MOI using serum-free Dulbecco’s modified Eagle’s medium (DMEM, VWR 45000-304) supplemented with 1x pen/strep, 1% nonessential amino acids, 2.5% HEPES, and cells were infected for one hour at 37°C. Cells were washed twice with sterile PBS solution (FisherScientific, SH30264FS), and appropriate culture medium was added for the duration of the experiment.

### Tissue Culture Infectious Dose 50 Assay

Supernatants from VSV-GFP, MERS-CoV and SARS-CoV-2 infected cells were titrated in triplicate on Vero76 cells using TCID_50_ assay. Briefly, 3×10^4^ cells were seeded per well of a 96-well plate. Following 24 hours of seeding, media was removed from the cells and 50 uL of serially diluted virus-containing supernatant was added to each well for 1 hour at 37°C. Following inoculation, the supernatant was removed and replaced with DMEM containing 2% FBS. The plates were incubated for one (VSV-GFP) to five days (MERS-CoV, SARS-CoV-2), and cytopathic effect was observed under a light microscope. Tissue culture infectious dose _50_/mL was calculated using the Spearman and Karber algorithm.

### Immunofluorescence assays

SEOV-infected cells were incubated at 37°C for 24 – 96 hours. At each time point, cell monolayers were washed twice with sterile PBS solution and fixed with ice cold 95% ethanol:5% acetic acid solution for 10 minutes on ice as previously described [45]. Cells were then washed twice with PBS and blocked with 3% fetal calf serum in PBS for 1 hr. Cells were probed overnight at 4°C for SEOV N protein (custom, Genscript) at dilution 1:400 in 1xPBS and with secondary antibody AlexaFluor 555 goat α mouse (Thermo Fisher Scientific, A-551 31570) at dilution 1:400 for two hours at room temperature. Nuclear stain DAPI was used at 1:1000 for ten minutes at room temperature (SeraCare, 5930-0006). Representative images were acquired on the EVOS FL Auto imaging system (Thermo Fisher Scientific, AMF7000).

### Immunoblots

Protein lysates were collected in 1X sample buffer and boiled for 10 minutes at 96°C. Proteins were separated by SDS-PAGE using homemade 10% polyacrylamide gels and semi-dry transferred to a 0.2 µm nitrocellulose membrane using the Trans-Blot Turbo Transfer System (BioRad, #1704270). Membranes were blocked using intercept blocking solution (Licor Biosciences, #927-60001) and probed with primary antibodies diluted in 50% intercept blocking solution and 50% TBS overnight on a rocker. Primary antibodies include VSV-M (Kerafast, #EB0011), MERS-CoV N (Sino Biological, #40068-RP01-200), IFIT1 (Thermo Scientific, #PA531254), GAPDH (EMD Millipore, #MAB374), and/or ACTB (Abcam, #ab8227). The appropriate secondary antibody (Licor Biosciences, #926-6870 or #926-32213) diluted in 50% intercept blocking solution and 50% TBS was then added for 60 minutes. Membranes were imaged and analyzed using the Odyssey imager (Licor Biosciences) and Image Studio Software (Licor Biosciences).

### RNA and qRT-PCR assays

Cells were lysed for RNA analysis using TRIzol Reagent (ThermoFisher Scientific, 15596026). RNA used for ANDV viral RNA quantification was purified using Zymo Research Direct-zol RNA Miniprep Plus kit (VWR 76211-340) according to manufacturer’s instructions. RNA concentrations were quantified using a NanoDrop UV-Vis Spectrophotometer (ND-1000). cDNA was synthesized using Applied Biosystems High-Capacity cDNA Reverse Transcription Kit (FisherScientific, 43-688-14) by adding 400ng total RNA to the reaction mixture containing random hexamers following manufacturer guidelines. Quantitative real-time PCR was performed using the Taqman Fast Advanced Master Mix (Thermo Fisher, 4444556) and the following primer sets as described by Warner et al.: ANDV S129f – AAGGCAGTGGAGGTGGAC, ANDV S291r – CCCTGTTGGATCAACTGGTT, ANDV TM - 56-FAM/ACGGGCAGC/ZEN/TGTGTCTACATTGGA/3IABkFQ (PMID: 33627395). Standard curve was generated with plasmid DNA containing the coding region of the ANDV nucleoprotein.

RNA was extracted from cells which received 100ng of nonlabelled poly(I:C) following the manufacturer’s instructions for the RNeasy Plus Mini Kit (Qiagen, #74136). Four hundred nanograms of purified RNA was reverse transcribed into cDNA following the manufacturer’s instructions for the iScript gDNA Clear cDNA Synthesis Kit (Bio-Rad, #1725035). qRT-PCR was performed using the SsoAdvanced Universal SYBR Green Supermix (BioRad, #1725274) to assess *IFNβ, IFIT1* and *GAPDH* transcript levels. The following primer sets were designed for this analysis: CpIFNb_F – GCAGCCCTGGAGGAAATC, CpIFNb_R – CCAGGCACATCTGCTGTAC, CpIFIT1_F – CTGTGATGCTTTGCCGAACC, CpIFT1_R – GGTTGGGCCTTGCTGAAATG, CpGAPDH_F – GGAGCGAGATCCCACCAACAT, and CpGAPDH_R – TGGAGTTGTCATACTTGTCCTGA.

### Reagent availability

The following top-performing cell lines from this study are available through BEI Resources and distributed by ATCC: Bulk, non-clonal *C. perspicillata* kidney cells immortalized with SV-40 T-antigen (NR-59778) and clonal CKg9 cells (NR-59779). All other cell lines are available upon request.

## Supporting information

Supplemental figures, tables and references

## AUTHOR CONTRIBUTIONS

ML, SNS, AB, and RKJ designed the study. CP provided animal tissues. ML and SNS isolated primary cells. ML and SF developed immortal cells. ML and MM developed clonal cells. ML, VG, CW, NGP, MM, SF, DF, AMK performed experiments. ML, VG and RJK assembled figures. ML, VG, AB, RJK, SNS, AMK wrote the manuscript. All authors contributed to the text.

## ACKNOWLEDGMENTS

We would like to thank Dr. Jay Hooper at USAMRIID for generously providing orthohantavirus M segment genes and B. Haagmans and R. Fouchier, Erasmus Medical Center, for providing MERS-CoV (isolate hCoV-EMC/2012). Research reported in this publication was supported by the National Institute of Allergy and Infectious Diseases of the National Institutes of Health (NIAID/NIH) under Award Numbers 1R21AI169527 (M.L. and A.B.), R 21AI156482 and P20GM134974 (R.K.J.), and Natural Sciences and Engineering Research Council of Canada (NSERC) Discovery Grant (RGPIN-2022-03010) awarded to A.B. This work was partly supported by an NSF Biology Integration Institute grant under award numbers NSF DBI 2021909 and 2213854 (S.N.S). V.G. is supported by an NSERC scholarship (#569587-2022). VIDO receives operational funding from the Government of Saskatchewan through Innovation Saskatchewan and the Ministry of Agriculture, and from the Canada Foundation for Innovation through the Major Science Initiatives Fund.

