## Supplemental figures, tables and references for "Viral susceptibility and innate immune competency of *Carollia perspicillata* bat cells produced for virological studies"

### Supplemental Materials

**Supplemental Table 1.** Cell types established from different bat species. Databases include the American Type Culture Collection (ATCC; USA), Collection of Cell Lines in Veterinary Medicine (CCLV; Germany), and Kunming Cell Bank of Chinese Academy of Science (KCB; China). References are included for cell lines not available through a database.

| Species | Tissue | Primary or Immortalized | Immortalization Technique | Cell Line | Commercial Source / Database | Ref. |
|---|---|---|---|---|---|---|
| ***Artibeus jamaicensis*** | Embryonic liver | Primary | | AjLi | | [1] |
| | Embryonic kidney | Primary | | AjLKi | | [2] |
| | Embryonic intestine | Primary | | AjIn | | [2] |
| | Kidney | Immortalized | SV40T | AJi | | [3] |
| | Intestine | Primary | | Intestinal Organoids | | [4] |
| ***Aselliscus stoliczkanus*** | Skin | | | STB-S2 | KCB #5301MAM-KCB05010ZJ | |
| ***Carollia perspicillata*** | Trachea | Immortalized | SV40T | CarperAEC.B | | [5] |
| | Lung | Primary | | CpLu | | [6] |
| | Kidney | Immortalized | SV40T | CarNi/1 | | [7] |
| | Trachea | Primary | | Airway Organoids | | [8] |
| | Lung | Primary | | Airway Organoids | | [8] |
| ***Cynopterus sphinx*** | Lung | | | GsnFB-L2 | KCB #5301MAM-KCB0502LZJ | |
| | Skin | | | GsnFB-S1 | KCB #5301MAM-KCB0501SZJ | |
| ***Desmodus rotundus*** | Embryonic liver | Primary | | DsLi | | [1] |
| | Embryonic kidney | Primary | | DsKi | | [1] |
| | Embryonic lung | Immortalized | SV40T | FLuDero | | [9] |
| ***Eidolon helvum*** | Embryonic kidney | Immortalized | SV40T | EidNi/41 | | [10] |
| | Trachea | Immortalized | SV40T | EidheAEC.B | | [5] |

| | | | | | | |
|---|---|---|---|---|---|---|
| | Lung | Immortalized | SV40T | EidLu/20 | | [10, 11] |
| | Lung | Immortalized | SV40T | EidLu/43 | | [10, 12] |
| | Kidney | Immortalized | SV40T | ZFBS13-75A | | [13] |
| | Kidney | Immortalized | SV40T | ZFBK13-76E | | [14] |
| ***Eonycteris spelaea*** | Lung | | | DB-L1 | KCB #5301MAM-KCB0683LZJ | |
| | Muscle | | | DB-M1 | KCB #5301MAM-KCB0683MZJ | |
| | Trachea | Primary | | Airway Organoids | | [15] |
| ***Epomops buettikoferi*** | Kidney | Immortalized | SV40T | EpoNi/22 | | [16] |
| ***Epomophorus gambianus*** | Kidney | Immortalized | SV40T | ZFBK11-97 | | [17] |
| ***Eptesicus fuscus*** | Kidney | Immortalized | MyPVT | Efk3 | Kerafast #CVCL_GZ34 | [18] |
| ***Eptesicus nilssonii*** | Kidney | Primary | | HAMOI-EnK | | [19] |
| ***Eptesicus serotinus*** | Brain | | | FLG-ID | CCLV | [20] |
| | Brain | | | FLG-R | CCLV | [20] |
| | Kidney | | | FLN-R | CCLV | [20] |
| ***Hipposideros abae*** | Lung | Immortalized | SV40T | HipaLu/24 | | [21] |
| | Lung | Immortalized | SV40T | HipaLu/27 | | [21] |
| ***Hipposideros armiger*** | Skin | | | GRB-S1 | KCB #5301MAM-KCB04046ZJ | |
| ***Hipposideros caffer*** | Embryonic | Immortalized | SV40T | HipaEm/5 | | [21] |
| | Embryonic | Immortalized | SV40T | HipEm/28 | | [21] |
| ***Hypsignathus monstrosus*** | Embryonic kidney | Immortalized | SV40T | HypNi/1 | | [16] |
| | Kidney | Immortalized | SV40T | HypNi/21 | | [21] |
| | Lung | Immortalized | SV40T | HypLu/2 | | [22] |
| | Embryonic Lung | Immortalized | SV40T | HypLu/45 | | [2] |

| | | | | | | |
|---|---|---|---|---|---|---|
| ***Macrotus waterhousii*** | Heart | | | Mw1Ht | ATCC #CRL-6013 *discontinued* | |
| | Lung | | | Mw2Lu | ATCC #CRL-6014 *discontinued* | |
| ***Miniopterus fuliginosus*** | Kidney | Immortalized | SV40T | YubFKT1 | | [17] |
| | Kidney | Immortalized | SV40T | YubFKT2 | | [23] |
| ***Molossus sinaloae*** | Embryonic liver | Primary | | MsLi | | [1] |
| | Embryonic kidney | Primary | | MsKi | | [1] |
| | Embryonic intestine | Primary | | MsIn | | [1] |
| ***Myotis altarium*** | Skin | | | SM-S1 | KCB #5301MAM-KCB04041ZJ | |
| ***Myotis daubentonii*** | Brain | Immortalized | SV40T | MyDauBrain/48 | | [21] |
| | Intestine | Immortalized | SV40T | MyDauDa/46 | | [21] |
| | Lung | Immortalized | SV40T | MyDauLu/47 | | [16] |
| | Kidney | | | MyDauNi/2 | | [7] |
| ***Myotis davidii*** | Kidney | Primary | | MdKi | | [24] |
| ***Myotis myotis*** | Brain | Immortalized | SV40T | MmBr | | [25] |
| | Tonsil | Immortalized | SV40T | MmTo | | [25] |
| | Peritoneal cavity | Immortalized | SV40T | MmPca | | [25] |
| | Nasal epithelium | Immortalized | SV40T | MmNep | | [25] |
| | Nervus olfactorius | Immortalized | SV40T | MmNol | | [25] |
| | Tail | Induced pluripotent stem cells | | Mmy iPSCs | | [26] |
| | Skin | | | MMY-S2 | KCB #5301MAM-KCB04044ZJ | |
| ***Myotis schreibersii*** | Lymph node | Primary | | MsLn | | [27] |
| | Kidney | Primary | | MsKi | | [27] |
| | Kidney | Immortalized | SV40T | SuBK12-08 | | [17] |
| ***Myotis velifer*** | Interscapular tumor | Cancerous | | Mvi/It | ATCC #CRL-6012 *discontinued* | [28] |
| | Muscle | | | Mvi/Mu | ATCC #CRL-6011 *discontinued* | |
| ***Pipistrellus ceylonicus*** | Embryonic | Spontaneously Immortalized | | NIV-BtEPC | | [29] |

| | | | | | |
|---|---|---|---|---|---|
| ***Pipistrellus nathusii*** | Kidney | Primary | | bKEC | [30] |
| ***Pipistrellus pipistrellus*** | Kidney | Immortalized | SV40T | PipNi/1 | [7] |
| | Kidney | Immortalized | SV40T | PipNi/3 | [7] |
| | Kidney | Immortalized | SV40T | PipNi/4 | [21] |
| ***Pipistrellus subflavus*** | Lung | Immortalized | hTERT | PESU-B5L | [31] |
| ***Pteropus alecto*** | Aorta | Primary | | PaAo | [32] |
| | Bone marrow | Primary | | PaBm | |
| | Brain | Primary | | PaBr | |
| | | Immortalized | SV40T | PaBrT01-03 | |
| | | Immortalized | hTERT | PaBrH01-07 | |
| | Fetus | Primary | | PaFe | |
| | | Immortalized | SV40T | PaFeT01-10 | |
| | Embryonic Membrane | Primary | | PaFm | |
| | Heart | Primary | | PaHe | |
| | Kidney | Primary | | PaKi | |
| | | Immortalized | SV40T | PaKiT01-03 | |
| | Liver | Primary | | PaLi | |
| | Lymph node | Primary | | PaLn | |
| | Lung | Primary | | PaLu | |
| | | Immortalized | SV40T | PaLuT01-04 | |
| | Muscle | Primary | | PaMu | |
| | Pharynx | Primary | | PaPh | |
| | Placenta | Primary | | PaPl | |
| | Salivary gland | Primary | | PaSg | |
| | Small intestine | Primary | | PaSi | |
| | Skin | Primary | | PaSk | |
| | Spleen | Primary | | PaSp | |
| | Testes | Primary | | PaTe | |
| | Thymus | Primary | | PaTh | |
| | Uterus | Primary | | PaUt | |
| ***Pteropus dasymallus yayeyamae*** | Kidney | Immortalized | SV40T | FBKT1 | [33] |
| ***Pteropus giganteus*** | Spleen | Immortalized | SV40T | IndFSPT1 | [17] |
| ***Pteropus pselaphon*** | 5th finger of right wing and skin | Immortalized | CDK4, CYCLIN D1, and TERT | Bff-K4DT | [34] |
| ***Rhinolophus affinis*** | Fetus | Primary | | BEF | [35] |

| | | | | | | |
|---|---|---|---|---|---|---|
| ***Rhinolophus alycone*** | Lung | Immortalized | SV40T | RhiLu/1.1 | | [6] |
| | Kidney | Immortalized | SV40T | RhiNi/1.2 | | [3, 21] |
| | Brain | Immortalized | SV40T | RhiBrain/4p | | [21] |
| ***Rhinolophus euryale*** | Lung | Immortalized | SV40T | RhiEuLu | | [21] |
| ***Rhinolophus ferrumequinum*** | Kidney | Immortalized | SV40T | BKT1 | | [17] |
| | Lung | Immortalized | SV40T | RhiFeLu | | [21] |
| | Fetus | Induced pluripotent stem cells | | Rfe iPSCs | | [26] |
| | Pulmonary | Immortalized | SV40T | RfPT | | [36] |
| | Brian | Immortalized | SV40T | RfBT | | |
| | Heart | Immortalized | SV40T | RfHT | | |
| | Kidney | Immortalized | SV40T | RfKT | | |
| | Lung | Immortalized | | GHB-L3 | KCB #5301MAM-KCB0605LZJ | |
| | Ear skin | | | GHB-S3 | KCB #5301MAM-KCB0605SZJ | |
| ***Rhinolophus landeri*** | Lung | Immortalized | SV40T | RhiLu/1.1 | | [7] |
| | Kidney | Immortalized | SV40T | RIKd | | [21] |
| ***Rhinolophus lepidus*** | Kidney | Spontaneously Immortalized | | Rhileki | | [37] |
| ***Rhinolophus pusillus*** | Lung | | | LHB-L2 | KCB #5301MAM-KCB04052ZJ | |
| | Muscle | | | LHB-M4 | KCB #5301MAM-KCB05058ZJ | |
| | Skin | | | LHB-S3 | KCB #5301MAM-KCB05012ZJ | |
| ***Rhinolophus sinicus*** | Kidney | Immortalized | SV40T | RsKT | | [24] |
| | Splenocytes | Primary | | BS | | [35] |
| | Skin | | | CHB-S2 | KCB #5301MAM-KCB05024ZJ | |
| | Intestine | Primary | | Intestinal Organoids | | [38] |

| | | | | | | |
|---|---|---|---|---|---|---|
| *Rousettus ageyptiacus* | Kidney | Immortalized | SV40T | RoNi/7.1 | | [16] |
| | Body of fetus | Immortalized | Adenovirus | R06E | | [39] |
| | Head of fetus | Immortalized | Adenovirus | R05T | | [39] |
| | Vertebrate column of fetus | Immortalized | Adenovirus | R05R | | [39] |
| | Lung | Primary | | RALU | | [40] |
| | Kidney | Primary | | RaKSM | | [40] |
| | Kidney | Immortalized | SV40T | RaKSM-2.5i | | [3] |
| | Kidney | Immortalized | SV40T | ZFBK15-137RA | | [23] |
| | Endometrium | Immortalized | SV40T | RoEnd/4 | | [21] |
| | Lung | Primary | | Airway Organoids | | [41] |
| | Intestine | Primary | | Intestinal Organoids | | [41] |
| *Rousettus leschenaulti* | Kidney | Immortalized | SV40T | DemKT1 | | [17] |
| | Intestine | Primary | | Intestinal Organoids | | [42] |
| | Lung | | | FFB-L3 | KCB #5301MAM-KCB0520LZJ | |
| | Skin | | | FFB-S3 | KCB #5301MAM-KCB0521SZJ | |
| *Tadarida brasiliensis* | Lung | Primary | | Tb1-Lu | ATCC #CCL-88 | |

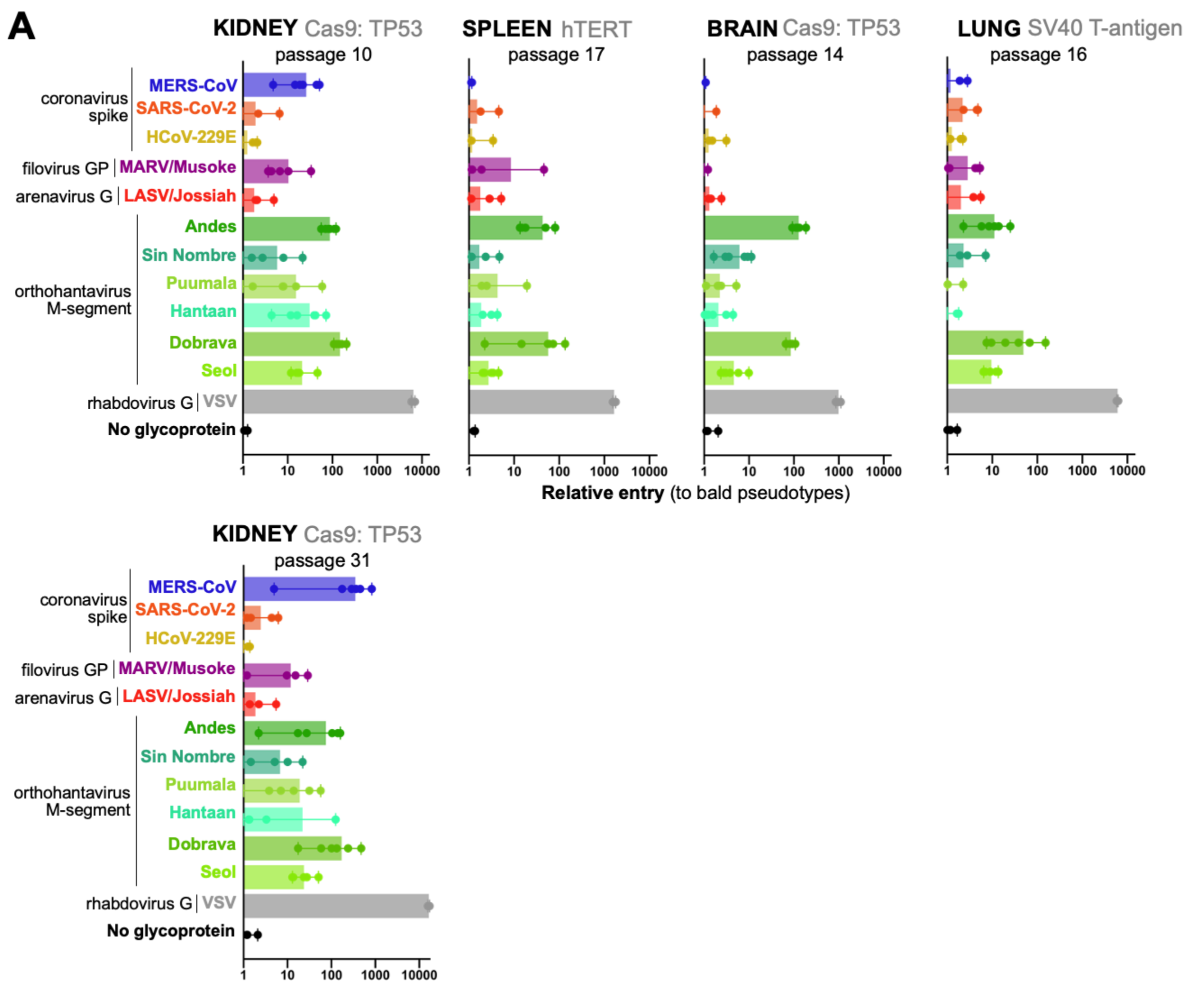


**Supplemental figure 2. Susceptibility of late-passage immortalized cultures. (A)** Immortalized *Carollia perspicillata* cells were maintained in culture for multiple passages and infected with the indicated pseudotypes.

#### SUPPLEMENTAL REFEREENCES


1. Moreira-Soto, A., C. Soto-Garita, and E. Corrales-Aguilar, *Neotropical primary bat cell lines show restricted dengue virus replication.* Comparative Immunology, Microbiology and Infectious Diseases, 2017. **50**: p. 101-105.
2. Miller, M.R., et al., *Broad and temperature independent replication potential of filoviruses on cells derived from old and new world bat species.* The Journal of Infectious Diseases, 2016. **214**(suppl_3): p. S297-S302.
3. Letko, M., A. Marzi, and V. Munster, *Functional assessment of cell entry and receptor usage for SARS-CoV-2 and other lineage B betacoronaviruses.* Nature microbiology, 2020. **5**(4): p. 562-569.
4. Hashimi, M., et al., *Antiviral responses in a Jamaican fruit bat intestinal organoid model of SARS-CoV-2 infection.* Nature Communications, 2023. **14**(1): p. 6882.
5. Eckerle, I., et al., *Bat airway epithelial cells: a novel tool for the study of zoonotic viruses.* PloS one, 2014. **9**(1): p. e84679.
6. Hoffmann, M., et al., *Differential sensitivity of bat cells to infection by enveloped RNA viruses: coronaviruses, paramyxoviruses, filoviruses, and influenza viruses.* PloS one, 2013. **8**(8): p. e72942.
7. Müller, M.A., et al., *Human coronavirus EMC does not require the SARS-coronavirus receptor and maintains broad replicative capability in mammalian cell lines.* MBio, 2012. **3**(6): p. 10.1128/mbio. 00515-12.
8. Su, A., et al., *Infection Studies with Airway Organoids from Carollia perspicillata Indicate That the Respiratory Epithelium Is Not a Barrier for Interspecies Transmission of Influenza Viruses.* Microbiology spectrum, 2023. **11**(2): p. e03098-22.
9. Sarkis, S., et al., *Development of molecular and cellular tools to decipher the type I IFN pathway of the common vampire bat.* Developmental & Comparative Immunology, 2018. **81**: p. 1-7.
10. Biesold, S.E., et al., *Type I interferon reaction to viral infection in interferon-competent, immortalized cell lines from the African fruit bat Eidolon helvum.* PloS one, 2011. **6**(11): p. e28131.
11. Ng, M., et al., *Filovirus receptor NPC1 contributes to species-specific patterns of ebolavirus susceptibility in bats.* Elife, 2015. **4**: p. e11785.
12. Hoffmann, M., et al., *Tetherin inhibits Nipah virus but not Ebola virus replication in fruit bat cells.* Journal of virology, 2019. **93**(3): p. 10.1128/jvi. 01821-18.
13. Maruyama, J., et al., *Characterization of the glycoproteins of bat-derived influenza viruses.* Virology, 2016. **488**: p. 43-50.
14. Ogawa, H., et al., *Characterization of a novel bat adenovirus isolated from straw-colored fruit bat (Eidolon helvum).* Viruses, 2017. **9**(12): p. 371.
15. Chan, L.L., et al., *Generation of self-replicating airway organoids from the cave nectar bat Eonycteris spelaea as a model system for studying host–pathogen interactions in the bat airway epithelium.* Emerging Microbes & Infections, 2023. **12**(1): p. e2148561.
16. Kühl, A., et al., *Comparative analysis of Ebola virus glycoprotein interactions with human and bat cells.* The Journal of infectious diseases, 2011. **204**(suppl_3): p. S840-S849.
17. Maruyama, J., et al., *Characterization of the envelope glycoprotein of a novel filovirus, lloviu virus.* Journal of virology, 2014. **88**(1): p. 99-109.
18. Banerjee, A., et al., *Generation and Characterization of Eptesicus fuscus (Big brown bat) kidney cell lines immortalized using the Myotis polyomavirus large T-antigen.* Journal of virological methods, 2016. **237**: p. 166-173.
19. Horie, M., et al., *Establishment and characterization of a cell line derived from Eptesicus nilssonii.* Journal of Veterinary Medical Science, 2016. **78**(11): p. 1727-1729.
20. Richter, M., et al., *The viral envelope is not sufficient to transfer the unique broad cell tropism of Bungowannah virus to a related pestivirus.* Journal of General Virology, 2014. **95**(10): p. 2216-2222.
21. Gützkow, T., *Interspecies-transmission of animal coronaviruses*. 2013, Hannover, Tierärztliche Hochsch., Diss., 2013.
22. Krüger, N., et al., *Attachment protein G of an African bat henipavirus is differentially restricted in chiropteran and nonchiropteran cells.* Journal of virology, 2014. **88**(20): p. 11973-11980.
23. Sato, M., et al., *Generation of bat-derived influenza viruses and their reassortants.* Scientific reports, 2019. **9**(1): p. 1158.
